# Cell size sensing in animal cells coordinates anabolic growth rates with cell cycle progression to maintain uniformity of cell size

**DOI:** 10.1101/123851

**Authors:** Miriam B. Ginzberg, Nancy Chang, Ran Kafri, Marc W. Kirschner

## Abstract

The uniformity of cell size in healthy tissues suggests that control mechanisms might coordinate cell growth and division. We derived a method to assay whether growth rates of individual cells depend on cell size, by combining time-lapse microscopy and immunofluorescence to monitor how variance in cell size changes as cells grow. This analysis revealed two periods in the cell cycle when cell size variance decreases in a manner incompatible with unregulated growth, suggesting that cells sense their own size and adjust their growth rate to correct aberrations. Monitoring nuclear growth in live cells confirmed that these decreases in variance reflect a process that selectively inhibits the growth of large cells while accelerating growth of small cells. We also detected cell-size-dependent adjustments of G1 length, which further reduce variability. Combining our assays with chemical and genetic perturbations confirmed that cells employ two strategies, adjusting both cell cycle length and growth rate, to maintain the appropriate size.

## Introduction

Uniformity of cell size is a consistent feature of healthy tissues. While different cell types can differ greatly in size, cells within a given tissue tend to be strikingly similar (1). In some tissues, loss of cell size uniformity is a characteristic of malignancy(2). Such observations raise the intriguing question of whether there are dedicated mechanisms that restrict cell size to a specific range. Is size uniformity the product of cellular processes that monitor cell size and correct deviations from a target size(1,3)?

Studies of yeast have long postulated the existence of a cell size checkpoint in the S. *cerevisiae*(4,5). In contrast, studies of animal cell size have remained inconclusive as to whether animal cells employ any form of size sensing (1,3). As early as 1965, A. Zetterberg published evidence consistent with the existence of a cell size checkpoint in human fibroblasts(6,7). Later literature, however, has predominantly suggested the opposite view, i.e. that growth and cell cycle progression are not coordinated(8–10). Similarly, in 2013 our group developed a mathematical analysis (ergodic rate analysis, ERA) that predicted the existence of a size-discriminatory process that coincides with S phase entry(11). However, since predictions of ERA are contingent on several assumptions, these findings did not resolve the controversy surrounding this topic(1,3). Equally important, while ERA hinted at the existence of a process that is influenced by cell size, it provided little insight into the nature of that process. This ambiguity was reflected in a more recent review in *Cell* concluding that “There is little evidence that a cell directly senses or uses some type of ruler to measure its size”(3). While these long-lasting controversies reflect the interest in the subject of size sensing, they also highlight the conceptual challenge impeding this field of study: what are the experimental criteria for detecting cell size sensing?

While it is easy to envision experimental assays of cell size, assays of size *sensing* are more challenging to conceptualize. In this study, we describe experimental approaches to assay size sensing by monitoring cell size variance. We also develop methods to separately assay the influence of cell size on cell cycle progression and on growth rate. With these approaches, we determine that animal cells monitor their size and correct deviations in cell size. Our results show that, like S. cerevisiae, animal cells that are smaller than their target size spend longer periods of growth in G1. Surprisingly, however, we found that in addition to a G1 length extension in small cells, animal cells also employ a conceptually different strategy of size correction. During two distinct points in cell cycle, anabolic growth rates are transiently adjusted so that small cells grow faster and large cells grow slower. These periods of growth rate adjustments function to lower cell size variability, promoting size uniformity. To our knowledge, such a reciprocal coordination of growth rate with cell size has not been previously observed in any organism.

## Results

### A size threshold regulates S-phase entry in animal cells

Previous literature has defined cell size as cell volume(12,13) or cell mass(14). In this study, we define cell size by a cell’s total macromolecular protein mass, as this metric most closely reflects the sum of anabolic processes typically associated with cell growth(15) and with activity in growth promoting pathways such as mTOR(16). In contrast, cell volume is a more labile phenotype, sensitive to ion channel regulation and fluctuations in extracellular osmolarity.

If cell size checkpoints do not exist, and the size of a cell is not a determinant of S phase entry, the reason that S phase cells are larger than G1 cells must be that S-phase cells have had more time to grow since their last division. Thus, to test whether a cell size checkpoint regulates S phase entry we compared cells in S-phase to cells of the same age (i.e. time elapsed since last division) that are still in G1.

To measure cell size together with the amount of time elapsed since a cell’s last division, we took the following approach. We used time-lapse microscopy to image live HeLa and Rpe1 cells for 1-3 days, and then immediately fixed them and applied AlexaFluor 647-Succinimidyl Ester (SE-A647), a quantitative protein stain that we previously established as a an accurate measure of cell mass (11). (For more supporting evidence on the use of this method, see supplementary file 1-1.) We computationally tracked thousands of individual cells over the course of the time-lapse movies and recorded the amount of time that elapsed between each cell’s “birth” via mitosis and its fixation, which we will refer to as the cell’s “age”. To distinguish S-phase cells from cells in G1, we used cell lines stably expressing mAG-hGem(17), a fluorescent reporter of APC activation and G1 exit. This experimental design allowed us to measure the age, size, and cell cycle stage of each cell at the time of fixation.

Figure 1A shows the average cell size as a function cell age (i.e. time since division). As expected, the average size of the oldest cells is double that of the youngest. The transient slowing of cell growth about 10 hours after birth (the average age of S-phase entry) is consistent with a behavior we and others(11,18) previously identified. In figure 1B, cell size vs. age (time since division) is plotted separately for G1 and post-G1 cells. As expected, cells in the first hours post-“birth” appear exclusively in the G1 stage, while later time points are characterized by heterogeneity in cell cycle stage, with some cells still in G1 while others have transitioned into S phase. Consistent with the existence of a cell size checkpoint, Fig 1B shows that S phase cells are significantly larger than G1 cells of the same age, even though both have been growing for the same amount of time. This result suggests that cells exit G1 in a size-dependent manner. Furthermore, the mean size of G1 cells plateaus as cells begins to enter S-phase, consistent with a mechanism where cells enter S-phase upon reaching a particular size threshold. The size difference between identically aged G1 and S phase cells is statistically significant (student’s t-test p< 3.92e-04) and was observed in three additional independent replicate experiments (p<0.009, p<5.80e-08, p<9.07e-15).

**Figure 1.**
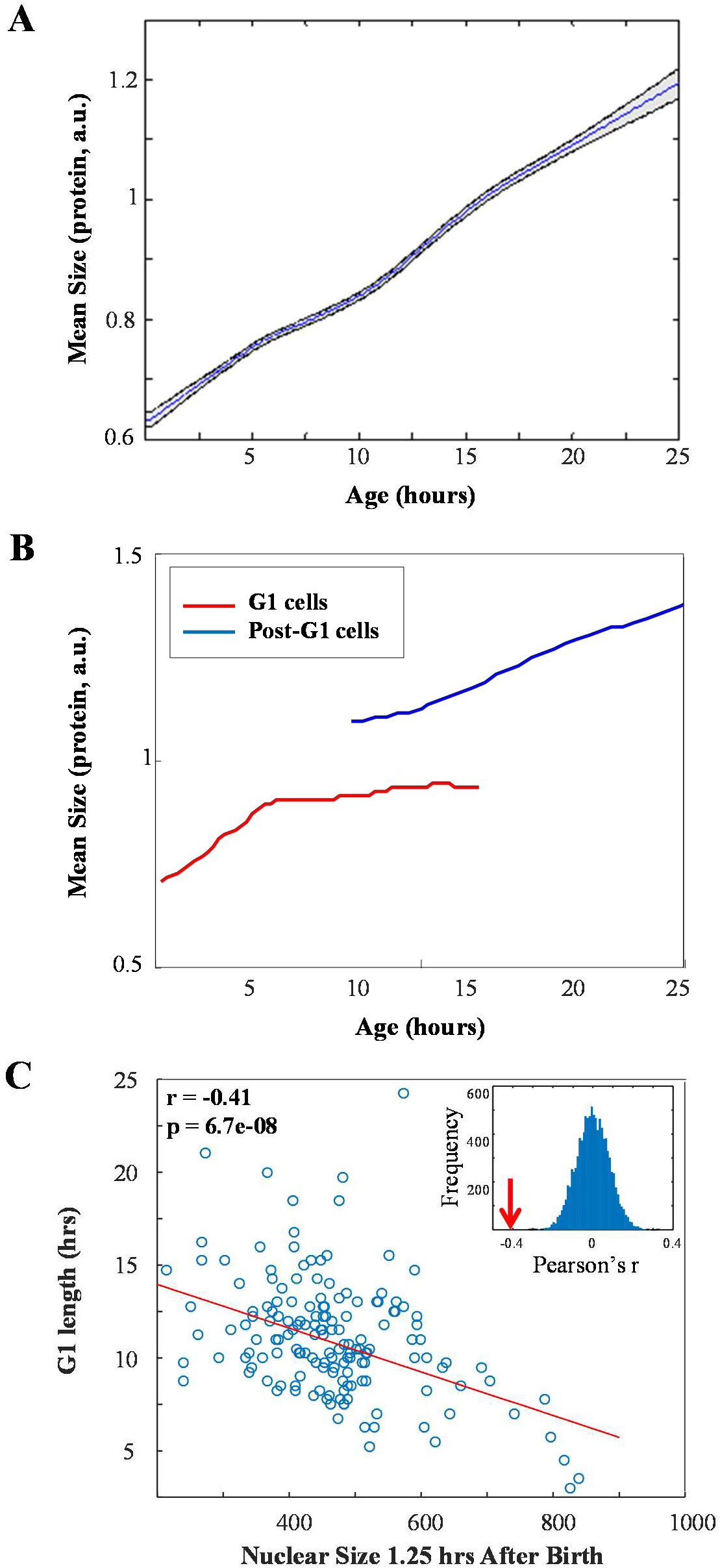
Small cells spend more time in G1. (A) Mean cell size as a function of age in HeLa cells. Shaded region marks 20% and 80% bootstrapping confidence intervals. (B) Mean size of G1 (red) and post-G1 (blue) HeLa cells as a function of cell age. Post-G1 cells are larger than G1 cells of the same age (student’s t-test p< 3.92e-04). Data shown in (A) and (B) is a representative example of four biological replicates (full experimental repeats), n=5537 cells. (C) Size (area covered in widefield fluorescence image) of HeLa nucleus 1.25 hours after birth vs. G1 duration. Line shows least-squares linear fit. Pearson’s r=-0.41, p=6.7x10^−8^, by two-tailed Student’s t-test. Inset shows distribution of Pearson’s r-values generated by randomizing the data, with arrow marking the r-value of non-randomized data. Data shown in (C) is a representative example of four biological replicates, n=158 cells.

If cells enter S-phase only after attaining a threshold size, cells that are born small are expected to have longer G1 periods. To test this, we assayed the correlation of cell size at birth with a direct measurement of G1-length in single cells. Since we cannot use the protein staining technique in live cells, we measured the size of the nucleus as a proxy for cell size, using time-lapse microscopy. In yeast, the nucleus is known to grow continuously throughout all stages of cell cycle and is correlated with cell size(19), and we found that the same is true in our experimental model (supplementary file 1-2). To further verify that nuclear growth accompanies cellular growth, independent of cell cycle progression, we arrested the cell cycle with aphidicolin and still observed a dramatic increase in nuclear size as cells grew (supplementary file 1-fig. S2C). When we monitored nuclear growth in single cells expressing nuclear-localized cell cycle markers(17), we observed a negative correlation between size of the nucleus at birth and G1 duration (fig. 1C). This result confirms that cells that are smaller at birth compensate with increased periods of growth in G1 and is consistent with the model of a cell size threshold at G1 exit.

To further test whether information about cell size is communicated to the cell cycle machinery, we examined the effect of slowing down cellular growth rates on the length of G1. If cells leave G1 only when a particular size has been reached, slowing down their growth rate is expected to prolong G1, as cells would require more time to reach the threshold size. Growth rate can be slowed down with drugs such as the mTORC1 inhibitor rapamycin. Culturing cells in 70 nM rapamycin caused a 60% decrease in the average growth rate and a compensatory 80% increase in G1 duration, resulting in only a 20% reduction in cell size (fig. 2A-E).

**Figure 2.**
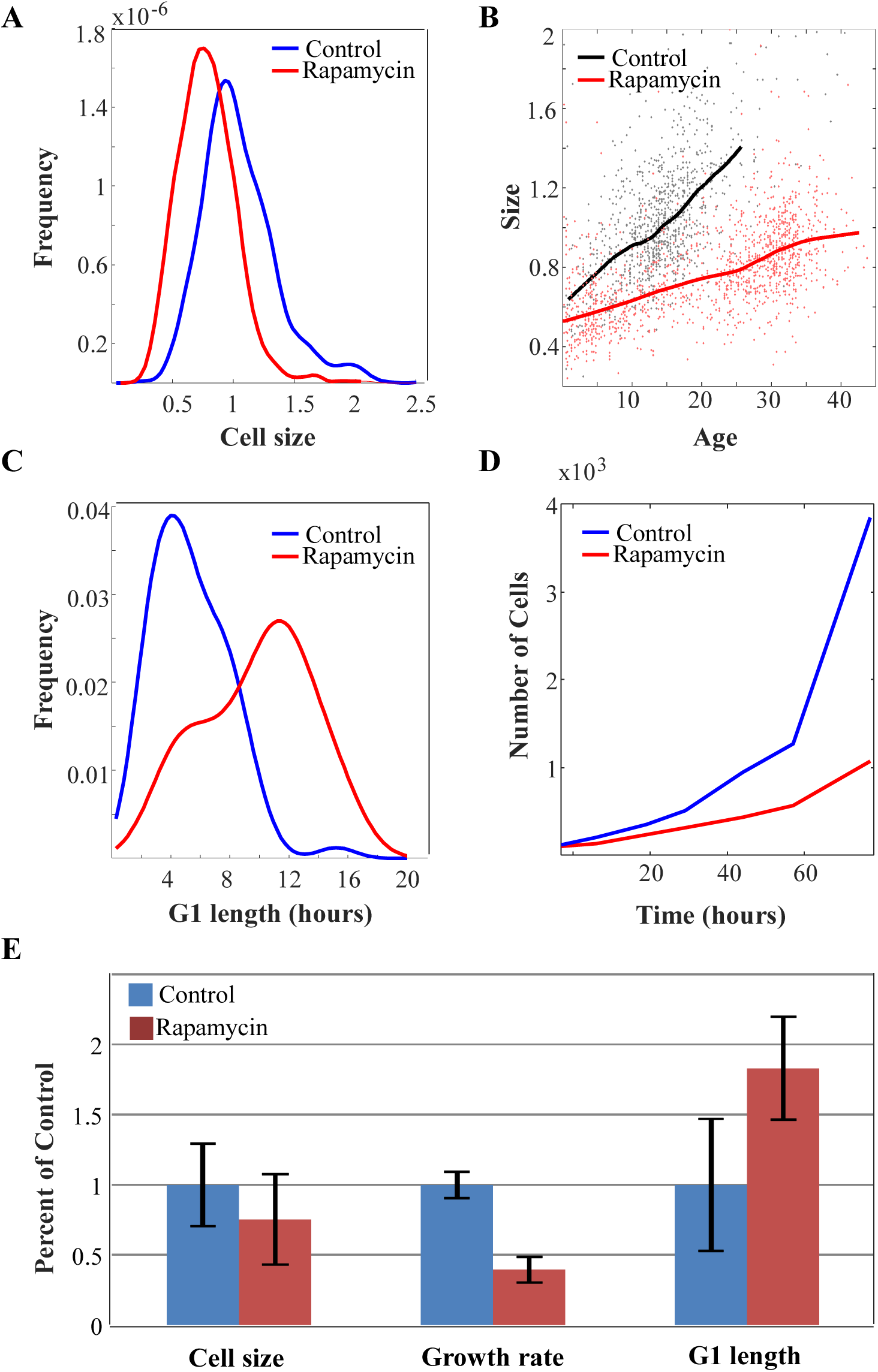
Slow growth rate in mTOR-inhibited cells is counteracted by increase in G1 duration. The effects of 70 nM rapamycin on the distribution of (A) cell size (protein content), (B) average growth rate, (C) G1-length, and (D) proliferation rate in Rpe1 cells. Solid lines in (B) show mean size vs. age for control and rapamycin-treated cells, while points represent single cell measurements. (E) Bar plot comparing the influence of rapamycin on growth rate, G1 length, and cell size. Bar heights represent mean, error bars represent standard deviation (n=975 cells in control, n=1268 cells in rapamycin). Data shown is a representative example of two biological replicates (full experimental repeats).

Figure 2 shows that the influence of rapamycin on growth rate and on G1 length is far more dramatic than its influence on cell size. Previous studies have already shown that in addition to its effect on cell size, rapamycin also alters G1 length(20–22). In those studies, the dual influence of rapamycin on both size and G1 length were interpreted to suggest two independent (pleiotropic) functions of the drug(23–25). However, our current results suggest an alternative, more parsimonious interpretation. Namely, that the influence of rapamycin on G1 length is an indirect consequence of its influence on growth rate. According to this interpretation, the inhibition of cell growth by rapamycin results in a gradual reduction of cell size which, in turn, triggers compensatory increases in G1. This new interpretation is consistent with the negative correlation that we observe between cell size and G1 length, even in the absence of rapamycin (fig. 1B,C). Additional support for this new interpretation will be provided in a later section of this study (fig. 5C,F), where we show that while the influence of rapamycin on growth rate is observed immediately after exposure of cells to the drug, the influence of rapamycin on G1 length is observed only after a significant delay. This suggests that G1 length extension is only indirectly related to the rapamycin treatment, and is mediated by some property that changes slowly over time, presumably cell size. Also in figure 5, we will show that this relationship of growth rate and cell size is persistent across a larger variety of pharmacological and genetic perturbations.

### The rate of cell growth is adjusted in a size-dependent manner

The results described above suggest that size uniformity of animal cells is maintained by keeping small cells in G1 for longer periods of growth. However, terminally differentiated animal cells do not divide and yet, often undergo precise adjustments of cell size(26). This suggests the presence of size specification mechanisms that are not dependent on regular cell divisions. Furthermore, the size threshold model is fundamentally limited in its ability to correct the size of very large cells. If some cells grow fast enough to double even in a very short cell cycle, regulation of cell cycle length alone cannot constrain size variability in the population.

The size of a cell is the product of both growth duration (cell cycle length) and growth rate. If cells have a mechanism to sense their own size, it may be that not only G1 length but also cellular growth rate is adjusted accordingly, to achieve the appropriate cell size. To investigate this possibility, we examined the relationship between cell size and growth rate.

If a cell can sense its own size and adjust its growth rate accordingly – just as a thermostat coordinates heat production with room temperature via short bursts of heat–we might expect these corrections to be transient and subtle. Experimental detection of transient changes in growth rates of single cells is difficult. To circumvent this difficulty, we derived an indirect inference method to assay whether growth rates of individual cells are dependent on cell size. Using the coupled measurements of cell size and cell age described above (fig. 1), we sorted cells based on their ages (i.e. time since division) and calculated the variance in cell size as a function of age.

In the absence of any size-dependent growth rate regulation, the variance in cell size is expected to increase with age, as cells grow, since individual cells in a population will vary in their growth rates. In any given time interval, fast-growing cells will accumulate more mass than slow-growing cells, thereby increasing disparities in cell size (fig. 3A). In fact, variance in size can only decrease if small cells grow faster than large cells (fig. 3B). This can be demonstrated by considering a time interval during which cells grow from *S*_1_ to *S*_2_. Cell size variance at any given time *t*_2_ is related to the cell size variance at any previous time *t*_1_ by:

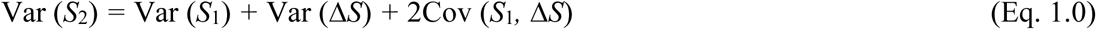

Therefore, the change in size variance follows:

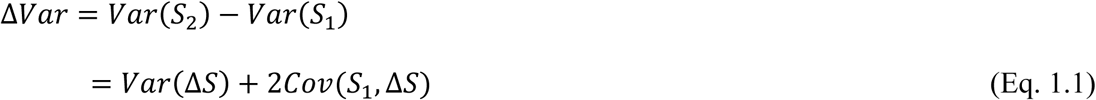

where ΔS is the change in cell size accruing over the time period Δ*t*, i.e. the growth rate. Since the variance in growth rates, Var (ΔS), is always positive (by definition), cell size variance can decrease with time (ΔVar <0) if (and only if) the correlation of cell size and growth rate is negative, i.e. Cov(S_1_, ΔS) < 0. This mathematical relationship can be exploited to experimentally detect periods of size-dependent growth rate regulation without directly measuring growth rates.

**Figure 3.**
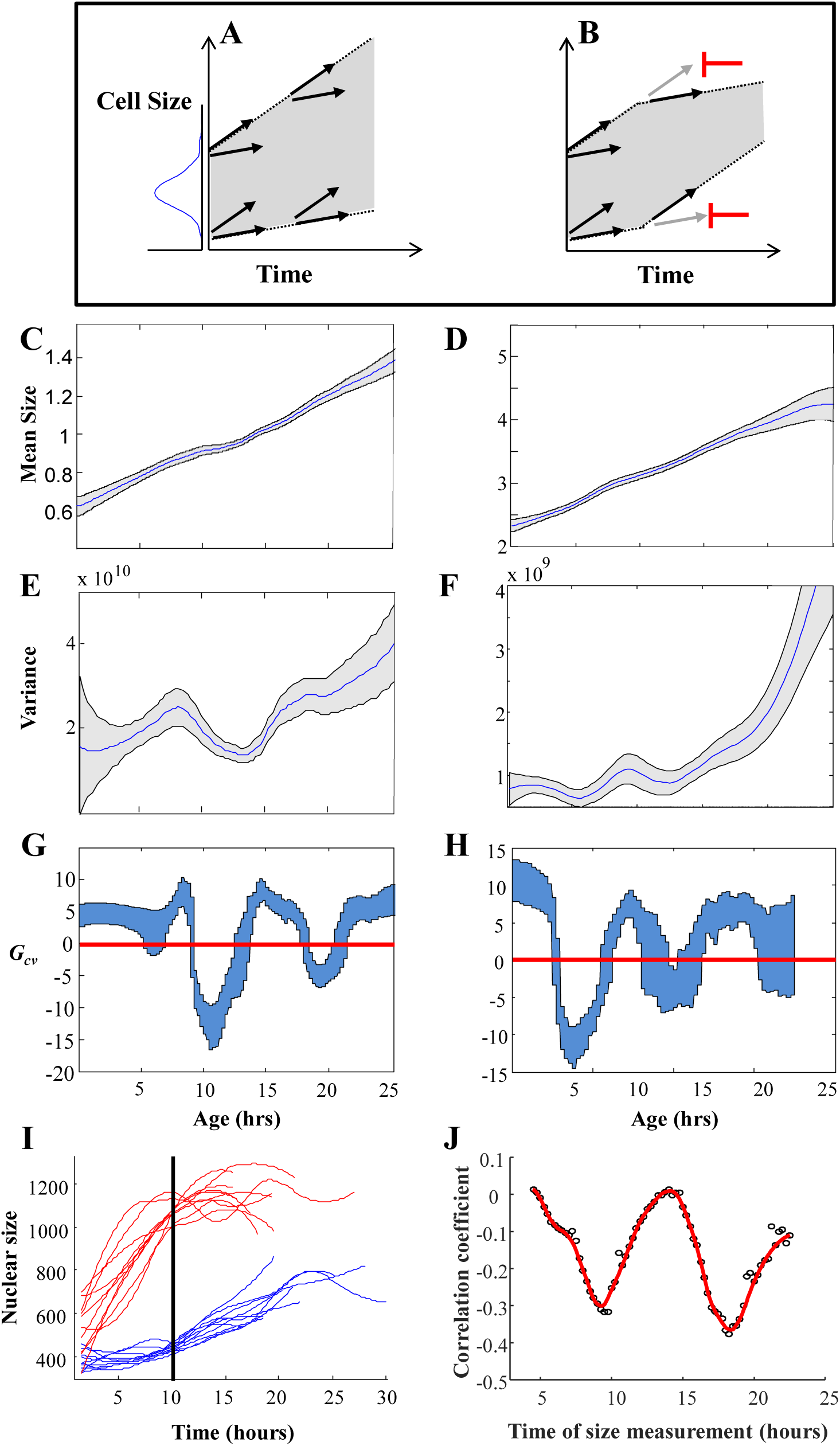
Cells adjust their growth rates in a size-dependent manner. (A) Variation in growth rates drives an increase in cell size variance over time. (B) Regulation that limits cell size variance must function via cross-talk between growth and cell size in individual cells. (C-H) Mean size, variance, and *G_cv_* plotted as a function of age for Rpe1 (C,E,G) and HeLa (D,F,H) cells. Shaded regions mark 20% and 80% bootstrapping confidence intervals. For Rpe1 plots (C,E,G), n=975 cells, and data shown is a representative example of four biological replicates. For HeLa plots (D,F,H), n=721 cells, and data shown is a representative example of four biological replicates. (I) Comparison of growth trajectories of smallest and largest cells. HeLa cells were sorted based on their size at 10 hours after birth. Nuclear size trajectories for the ten largest (red) and smallest (blue) cells are shown. (J) Correlation (Pearson’s r) between size and subsequent growth rate (2.5 hours later), calculated as a function of age (time since birth). Data shown in (I) and (J) is a representative example of four biological replicates, n=158 cells.

Figure 3 shows that periods of decreasing cell size variance do, in fact, occur during the cell cycle of both HeLa and RPE1 (Fig. 3C-F). During these periods, cells grow (mean size increases), and yet become more similar in size. This indicates that cellular growth rates are regulated to correct aberrant cell sizes. To quantify the strength of this regulation, we normalized the change in size variance, Δ*Var* = *Var*(*S_2_*) *– Var*(*S_1_*), by the amount of growth that has occurred (i.e. the change in mean size, 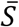) to define the *coefficient of growth rate variation*, designated *G_cv_*.

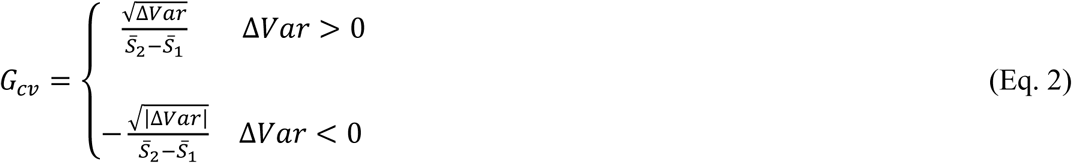

The *G*_*cv*_-value analysis is a new method to interrogate size control in growing cells. As long as the mean cell size is increasing over time, *G_cv_* can be interpreted as follows. If a cell’s growth rate is independent of its size, *G_cv_* directly equals the coefficient of variation (CV) of cellular growth rates. This can be shown by substituting the relationship in eq. 1 into the formula for *G*_*cv*_, noting that, if growth rates are size-independent, variance will increase over time and 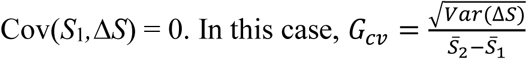.

Negative values of G_cv_ (arising from dips in variance) imply that growth rate is actually not independent of size, and that cell size and subsequent growth rate are negatively correlated. We can also note that if *G_cv_* is much higher than is plausible for the CV of growth rates (which will equal 1 if growth is a Poisson process), it is likely that growth is positively correlated with size, as in the case of exponential growth.

Plotting *G_cv_* versus cell age consistently reveals two distinct periods during which cell size and growth rate are negatively correlated (i.e. *G_cv_* is negative), in both HeLa and Rpe1 cells (Fig. 3G,H). This result is striking, because it suggests communication between cell size and cellular growth rate that is transiently established twice during the cell cycle, perhaps due to cell-cycle-dependent signalling linking size and growth rates.

The analysis presented in figure 3 does not discriminate G1 cells from S phase cells and, consequently, the observed drops in variance cannot be explained by a cell-size checkpoint gating G1. Therefore, figure 3 clearly reveals two distinct times in the cell cycle, where the growth rate of individual cells is selectively repressed in large cells or accelerated in small cells – increasing uniformity of cell size. The only alternative possibility is that cells are removed from the distribution, by either dividing or dying in a size-dependent manner. This is very unlikely to be the case, since the dips occur earlier than cells start dividing. (This is true for at least the first dip in Rpe1 and both dips in HeLa, where the mean cell cycle length is 21 hrs, with a standard deviation of 1.6 hrs. See supplementary file 1-3 for a sample cell cycle length distribution.) There was also a very low rate of apoptosis (<1% of cells imaged from birth that died before dividing).

The results of the *G_cv_*-value analysis are surprising. G1 checkpoints that link cell cycle progression to cell size have been previously observed in yeast(4). A few reports have suggested that G1 duration depends on growth rate rather than cell size(27,28). However, a process that stabilizes cell size by transient corrections of growth rate has not been previously observed in any organism. In fact, such a possibility was never previously hypothesized except in our own previous publication of ERA(11), but since ERA is fundamentally incapable of distinguishing changes in growth rate from changes in cell cycle progression rate, that suggestion remained a speculation. As such, this result represents the first clear evidence that cellular growth rates are adjusted in a cell-size-dependent manner to increase uniformity in a population.

To test the conclusions of the *G_cv_*-value analysis, we asked whether the correlation of growth rate and cell size could be observed directly in live cells. Using time-lapse microscopy, we monitored nuclear growth in hundreds of live cells over several days. Comparing growth trajectories collected from the largest and smallest cells in the population provided a dramatic demonstration that growth rates are, indeed, reciprocally coordinated with cell size (fig. 3I). Because growth rates became slower in large cells and faster in small cells, individual cells that were initially quite different in size gradually become more uniform in size, explaining the decreases in cell size variance shown in figure 3E-H. Furthermore, as predicted by the *G_cv_*-value analysis, the negative correlation of cell size and subsequent growth rate was not continuously present, but was transiently established twice during the cell cycle (fig. 3J).

### Growth rate and cell cycle length are coordinated to maintain cell size at fixed values

The *G_cv_*-value analysis and live-cell tracking of nucleus size all indicate that cellular growth rates are regulated in a manner that reduces cell size variability. Taken together, our results suggest that cells employ two separate strategies to correct deviations from their appropriate size: (1) small cells spend more time in G1 and (2) small cells grow faster than large cells (fig. 4). A prediction of this dual-mechanism model is that perturbations that slow the cell cycle would be counteracted by a compensatory decrease in growth rate, allowing cells to accumulate the same amount of mass despite the longer periods of growth. Conversely, perturbations that reduce growth rate would be counteracted by a compensatory lengthening of G1. To experimentally lengthen the cell cycle, we treated cells with a variety of pharmacological inhibitors of the cyclin dependent kinases or cdc25. Drugs were used at concentrations that slow the proliferation rate but do not cause cell cycle arrest (supplementary file 1-4). To experimentally decrease growth rates, we used cycloheximide, as well as the mTOR inhibitors Torin-2 and rapamycin, also at doses where cells maintain viability and proliferation (supplementary file 1-4). We then monitored growth and proliferation over time, by fixing samples at intervals and measuring the bulk protein content (total SE-A647 intensity of sample) and number of cells present. From these measurements, we calculated the average growth rate and cell cycle length of cells in each condition. Cell cycle length was also independently measured, by monitoring the proliferation of live cells in each condition with differential phase-contrast microscopy over the course of three days.

**Figure 4.**
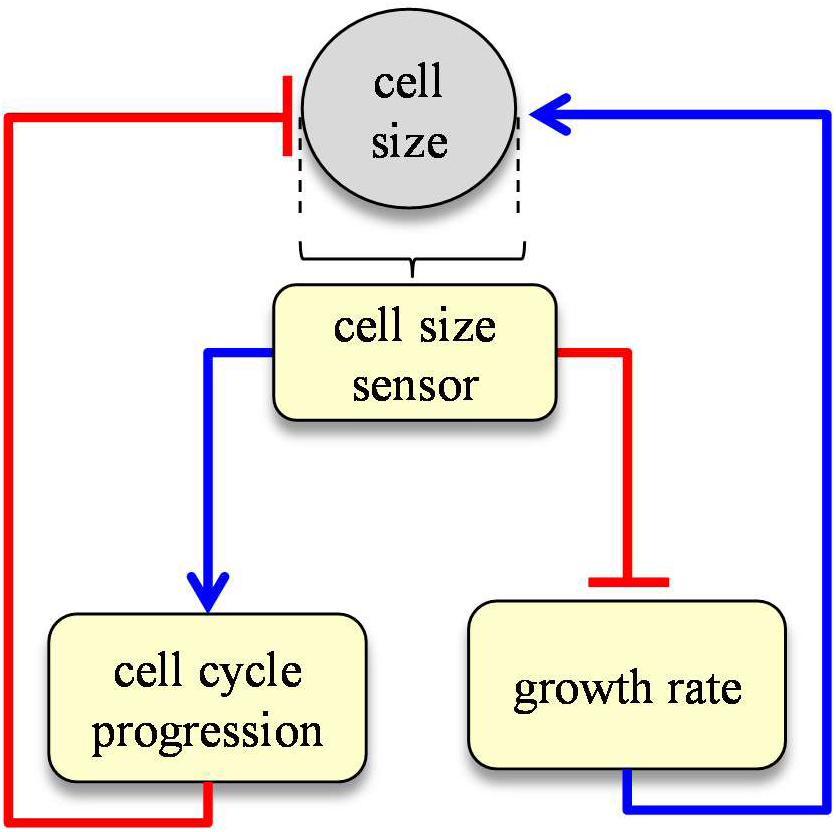
Dual-mechanism model of cell size specification. Data presented here is consistent with a model where cells sense their own size and employ two strategies, adjusting both growth rate and cell cycle length, to correct aberrant sizes.

Figure 5A,B shows that chemical perturbations of growth rate and perturbations of cell cycle length had surprisingly small effects on cell size. Consistent with the dual-mechanism model, treatment with all but one of the tested compounds resulted in coordinated changes of both cell cycle length and growth rate, such that normal cell size was maintained (fig. 5C, fig. 6C,D,F,G). Furthermore, while the inhibitors of cell cycle regulators produced an immediate effect on cell cycle length, their effect on growth rate was observed only after prolonged treatment (Fig. 5E). This suggests that the effect of these inhibitors on growth rate is indirect and is mediated by a property that accumulates over time, presumably cell size. Similarly, cycloheximide, rapamycin and Torin-2 induced an immediate decrease in growth rate, while the effect of these drugs on cell cycle length was observed after a delay (Fig. 5F). Naively, changing how fast cell grow or changing the amount of time cells grow should result in proportional changes to cell size. These results, however, demonstrate the surprising stability of cell size, in the face of perturbations to either growth rate or cell cycle progression, and reveal a compensatory relationship between these two processes.

**Figure 5.**
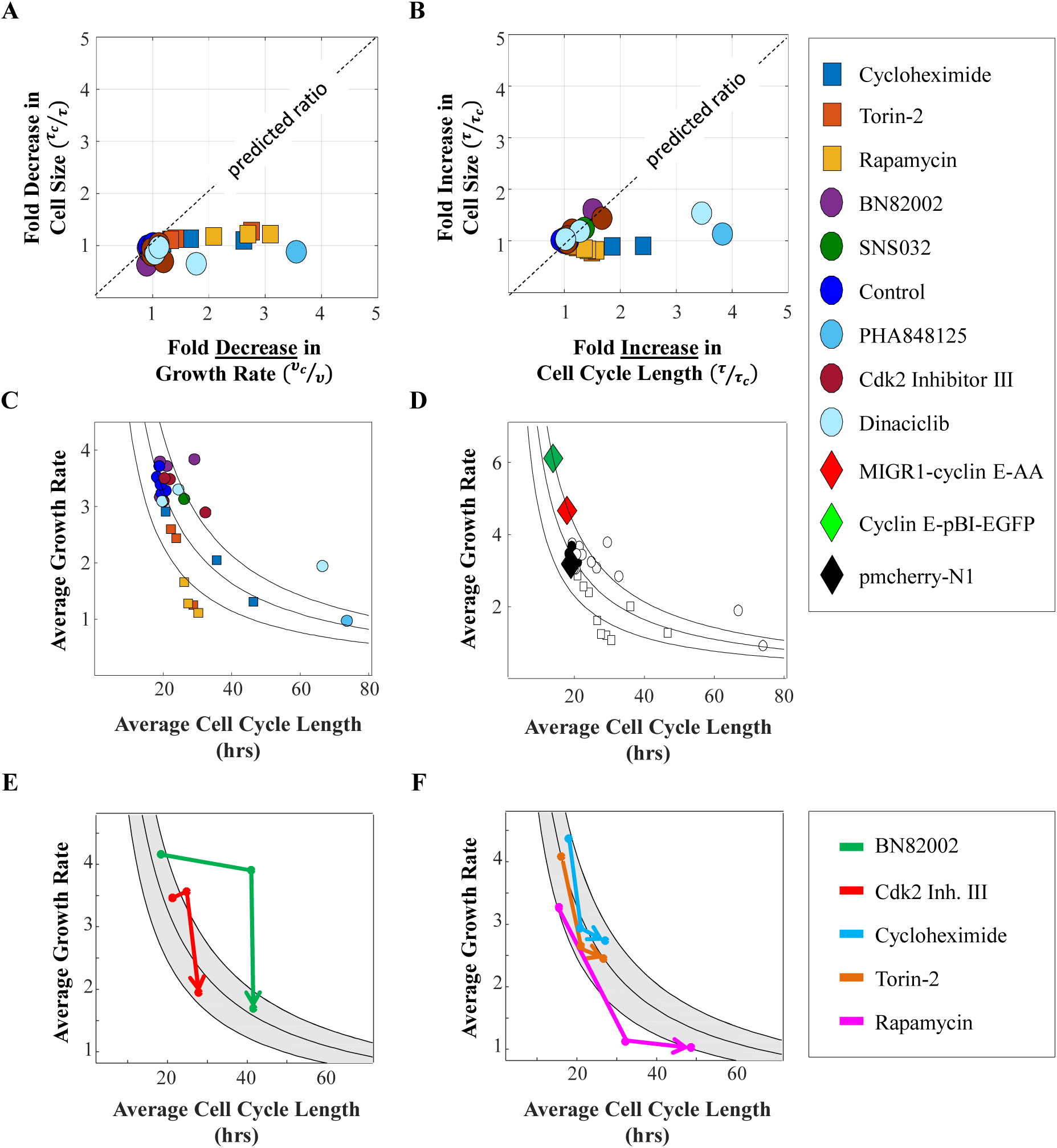
Compensatory relationship between cell cycle length and growth rate. (A-D) Cells were treated with varying doses of drugs inhibiting cell cycle progression (circles) or protein production (squares). (A) The fold change in cell size (after 44 hrs in drug) vs. fold change in growth rate, relative to untreated cells, is plotted for each condition. Dotted line denotes proportional relationship expected if there is no coordination between growth and cell cycle progression. (B) The fold change in cell size (after 44 hrs in drug) vs. fold change in cell cycle length, relative to untreated cells, is plotted for each condition. (C) Mean growth rate vs. mean cell cycle length are plotted for cells in each condition, calculated from measured increases in bulk protein and number of cells over the course of a 68-hour incubation. Black lines mark iso-cell-size contours. Region between lines spans a 25% shift in cell size. (D) Cells were transfected with plasmids encoding wild-type or degradation-resistant cyclin E under control of a constitutive promoter (diamonds). Mean growth rate vs. mean cell cycle length are plotted for cells in each condition, overlaid on the plot shown in (C). Calculation of the Pearson correlation coefficient between log(growth rate) and log(cell cycle length), using the data shown in (D), yields r = -0.76, p = 2.1x10^−7^, by two-tailed Student’s t-test, indicating a significant inverse correlation between growth rate and cell cycle length. (E,F) Mean growth rate vs. mean cell cycle length for cells treated with drugs inhibiting cell cycle progression (E) or protein production (F), calculated for three time intervals during drug treatment: 0-14 hrs, 14-44 hrs, 44-68 hrs. Lines connect time-points in sequential order, with arrowheads pointing to latest time-point. Data is shown for 25 uM BN82002 (E, green), 1.5 uM Cdk2 Inhibitor III (E, red), 0.06 uM Cycloheximide (F, blue), 10 nM Torin (F, orange), and 7 uM Rapamycin (F, purple). For (A-F), each treatment was done in duplicate, with several thousand cells in each sample.

**Figure 6.**
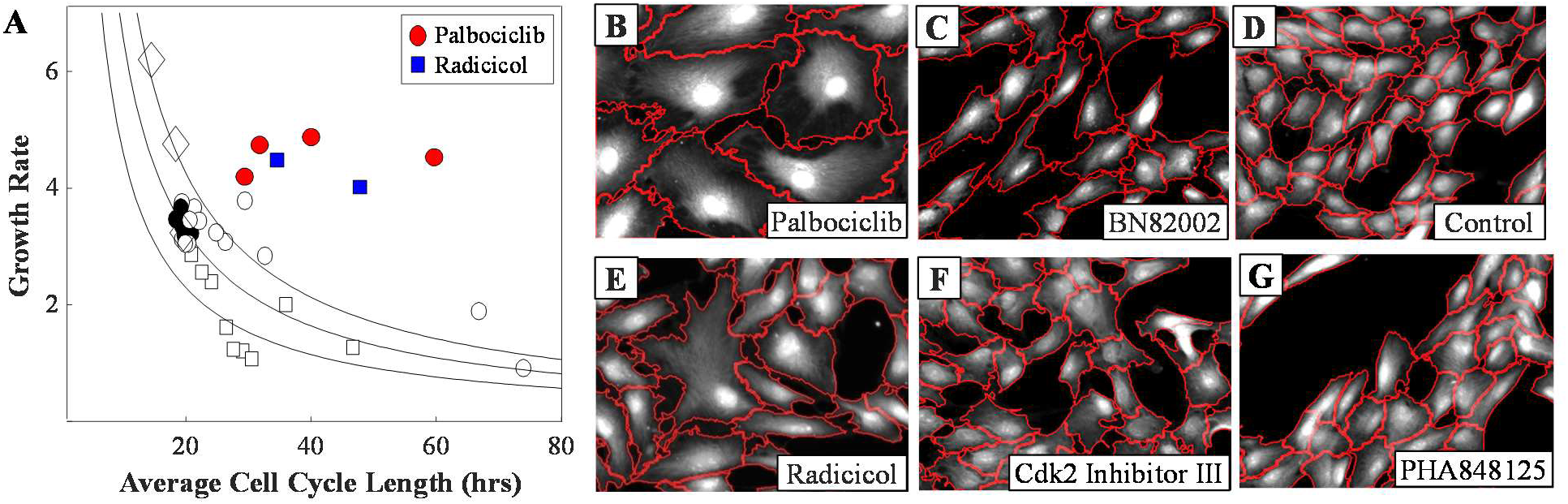
Perturbing the relationship between cell cycle length and growth rate. (A) Cells were treated with varying doses of palbociclib or radicicol. Mean growth rate vs. mean cell cycle length are plotted for cells in each condition, overlaid on the plot shown in figure 5D. (B-G) Representative images of cells treated with 500 nM palbociclib (B), 25 uM BN82002 (C), 250 nM radicicol (E), 5 uM Cdk2 Inhibitor III (F), 175 nM PHA848125 (G), and unperturbed cells (D). Red lines delineate cell boundaries.

To further test these conclusions, we overexpressed cyclin E to shorten the duration of cell cycle. Since CDK inhibitors that lengthened cell cycle were counteracted by a reduction in growth rate, we asked whether shortening the cell cycle would result in an increased growth rate. Figure 5D shows that, as anticipated, the shorter cell cycles caused by cyclin E overexpression are compensated for by faster rates of cell growth, rendering cell size relatively unchanged.

Of the tested perturbations to cell cycle regulators, only the cdk4/6 inhibitor palbociclib increased cell cycle length without a compensatory decrease in growth rate, causing an unusual increase in cell size (fig. 6A,B). Future studies investigating how palbociclib perturbs the crosstalk between cell cycle progression and cell growth may provide clues to the mechanisms of cell size sensing.

A quantitative prediction of the dual-mechanism model is that growth rate, *v*, and cell cycle length, τ, are coordinated to maintain cell size at its fixed target size:

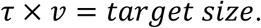

This means that growth rate and cell size are related by, 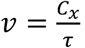, where *C_x_* is a constant target size. Figure 5 shows that our measurements are in agreement with this prediction. The curves in figure 5C-F are iso-cell-size contours and represent all paired values of growth rate and cell cycle length that correspond to one particular cell size. A notable feature of figure 5C,D is the significant influence of drugs on both growth rate and cell cycle length, in contrast to the relatively small influence of those same drugs on cell size. Naively, one would expect a drug that decreases growth rate by 50% would result in a 50% decrease in cell size. Results from our experiment show that, for many drugs, this expectation is false. While drug treatments produced up to fourfold changes in both growth rate and cell cycle length, the resulting changes in cell size were mostly under 30%. As mentioned above, while the influence of CDK inhibitors on cell cycle progression was immediate, their influence on growth rate was observed only after many hours of drug treatment (Fig. 5E), supporting the conclusion that the decrease in growth rate is a response to the gradual increase in cell size that occurs during a lengthened cell cycle. Conversely, while the influence of mTOR inhibitors on growth rate was immediate, their influence on cell cycle progression was observed only after a delay (Fig. 5F). These imply that cells have evolved regulation that buffers cell size from changes in cell cycle length or growth rate. In this light, it may be interesting to examine why cells have evolved to tolerate changes in cell cycle length and growth rate but not changes in cell size.

Finally, in an attempt to disrupt the processes that maintain cell size uniformity, we treated cells with the HSP90 inhibitor radicicol. HSP90 is known to suppress phenotypic variability(29) and has also been found to regulate both cell cycle progression(30,31) and mTORC1 mediated growth in response to cellular stress(32). Remarkably, culturing cells in radicicol not only decoupled growth rate and cell cycle length, but also resulted in a marked increase in cell size heterogeneity (fig. 6A,E). This finding illustrates that cell size uniformity is a regulated phenotype that can be perturbed by disrupting the coordination of growth and cell cycle progression.

## Discussion

The subject of cell size uniformity has been largely unexplored. It was not even clear whether size uniformity is important or why size is so uniform despite variation in growth rates and cell cycle lengths. In this study we show that cells that have escaped their appropriate size range are corrected by two separate mechanisms: regulation of G1 length and regulation of growth rate. To reveal these mechanisms, we circumvented the experimental difficulty of measuring cellular growth rates by developing methods that infer growth rate regulation from simpler measurements. With these approaches, we not only provide the first definitive proof that the size of an individual animal cell is autonomously monitored and regulated, but also provide accessible experimental assays for the study of growth rate regulation.

The stability of cell size in proliferating populations is a clear refutation of the notion that size is simply the quotient of growth and division rates, which are independently regulated with no feedback between them(10). Not only is there feedback, but it robustly maintains cell size despite strong perturbations of growth and cell cycle progression. The feedback that maintains size and size variation in growing cells suggests the presence of a size sensor. With an understanding that this sensor feeds back on both growth and cell cycle progression in opposite ways, and with assays that can separately quantify each mode of regulation, we are in a position to uncover its molecular identity.

In tissues, growth and shrinkage are often due to control of cell proliferation and apoptosis but some cell types such as neurons and muscle cells are controlled by cell growth rather than proliferation. Hence, in non-dividing cells that maintain size we may find part of the circuitry used in proliferating cells. Although pathways, such as mTOR, that promote cell growth have been delineated(16,33), it remains unclear how a common set of pathways may specify a different size in each cell type, while ensuring precise uniformity within cells of a common type. The approaches developed here can be combined with perturbations of these pathways to answer this fundamental question. With that knowledge, we can begin to explore the consequences of cell non-uniformity in normal cell function and particularly in development, disease, and aging.

## Methods

### Cell culture

Cells were grown in Dulbecco’s Modification of Eagles Medium (Cellgro, 10-017-CV), supplemented with 10% Fetal Bovine Serum (FBS, Gibco, 26140) and 5% Penicillin/Streptomycin (Cellgro 30-002-CI), incubated at 37°C with 5% CO2. All measurements were made when cells were 50-75% confluent, to avoid the effects of sparse or dense culture on cell growth and proliferation.

### Cell cycle markers

To distinguish S-phase cells from cells in G1, we used cell lines stably expressing nuclear-localized fluorescent cell cycle markers (A. Sakaue-Sawano *et al*., *Cell*. 132 (2008) 487–498).

### Time-lapse microscopy

Cells were seeded in 6-well, glass-bottom (No. 1.5) plates one day prior to imaging. Leibovitz’s L-15 Medium (ThermoFisher, 21083027) with 10% FBS was used during image acquisition, with a layer of mineral oil on top of the media to prevent evaporation. Microscope was fitted with an incubation chamber warmed to 37°C. Widefield fluorescence and phase contrast images were collected at 15 minute intervals on a Nikon Ti motorized inverted microscope, with a Nikon Plan Fluor 10x 0.3 NA objective lens and the Perfect Focus System for maintenance of focus over time. A Lumencor SOLA light engine was used for fluorescence illumination, and a Prior LED light source was used for transmitted light. A Prior Proscan III motorized stage was used to collect images at multiple positions in the plate during each interval. Images were acquired with a Hamamatsu ORCA-ER cooled-CCD camera controlled by MetaMorph software. After two days of time-lapse imaging, cells were fixed and stained as described below. Images of fixed cells were acquired on the same microscope, at the same stage positions, and an image-registration tool was designed in Matlab to match individual cells in the timelapse movies to the corresponding cells in the fixed images.

### Fixation and staining

Cells were fixed in 4% paraformaldehyde (Alfa Aesar, 30525-89-4) for 10 minutes, then permeabilized in chilled methanol for 5 minutes. Cells were stained with 0.04 ug/mL Alexa Fluor 647 carboxylic acid, succinimidyl ester (SE-A647, Invitrogen A-20006), to label protein. DNA was stained with DAPI (Sigma D8417).

### Pharmacological and genetic perturbations

To slow growth rate the following drug treatments were used: cycloheximide (1, 0.6, and 0.06 uM, Sigma C4859), Torin 2 (10, 5, and 2.5 nM, Tocris 4248), rapamycin (7, 0.7, and 0.07 uM, CalBiochem 553211). To slow the cell cycle, the following drug treatments were used: BN82002 (25, 12.5, 6.2, 3.1, 1.6, and 0.78 uM, Calbiochem 217691), SNS032 (39 and 9.8 nM, Selleckchem S1145), PHA848125 (175 nM, Selleckchem S2751), Cdk2 Inhibitor III (5, 1.5, 0.75, and 0.38 uM, Calbiochem 238803), Dinaciclib (10, 5, and 2.5 nM, Selleckchem S2768), Palbociclib (500, 250, 125, and 62.5 nM, Selleckchem S1116).

To promote cell cycle progression, cells were transfected with plasmids encoding wild-type or degradation-resistant cyclin E under control of a constitutive promoter. Cyclin E-pBI-EGFP was a gift from Bert Vogelstein (Addgene plasmid # 16654)( Rajagopalan, H. *et al. Nature* 428 (2004) 77–81). MIGR1-cyclin E-AA was a gift from Alex Minella (Addgene plasmid # 47498)( Xu, Y., Swartz, K. L., Siu, K. T., Bhattacharyya, M. & Minella, A. C. *Oncogene* 33 (2014) 3161–3171). Since the transfection process can slow proliferation, the growth rate and cell cycle length of cells overexpressing cyclin E were compared to those of cells transfected with a control plasmid, pmCherry-N1 (Clontech, 632523). For comparison with drug-treated cells (fig. 5D), growth rates and cell cycle lengths of transfected cells were normalized so that cells expressing pmCherry-N1 matched DMSO-treated cells (controls).

Radicicol (Tocris, 1589) treatments were done at concentrations of 250 and 500 nM. Cells were imaged using a Perkin Elmer Operetta high content microscope, controlled by Harmony software, with an incubated chamber kept at 37°C and 5% CO_2_ during live-cell imaging. A Xenon lamp was used for fluorescence illumination, and a 740 nm LED light source was used for transmitted light. To monitor proliferation of live cells, differential phase contrast images were collected every 6-12 hours, over the course of three days, using a 10x 0.4 NA objective lens. A plot of the number of cells vs. time was fitted to an exponential growth model to calculate the average cell cycle time.

Samples were fixed after 14, 44, and 68 hours in each drug. After fixation and staining with SE-A647 (protein) and DAPI, fluorescence images were collected with a 20x 0.75 NA objective lens. The bulk protein content (total SE-A647 intensity of sample) and number of cells were measured in each sample. From these measurements, we calculated the average growth rate and cell cycle length of cells in each condition, by fitting all data points (from two replicates of each condition) to an exponential growth model. Cell cycle length was also independently measured, by monitoring the proliferation of live cells in each condition with differential phase-contrast microscopy as described above.

### Image analysis and data analysis

All image analysis (cell segmentation, tracking, measurements of fluorescence intensity and nuclear size) and data analysis was performed with custom-written tools in Matlab. The mean and variance of cell size as a function of age (figure 3) were calculated by nonparametric regression, as described by Wasserman (34).

## Acknowledgements

Microscopy was done at the Nikon Imaging Center at Harvard Medical School, with the help of Jennifer Waters and Lara Petrak. Celina Qi assisted in computational analysis of time-lapse movies. We would like to thank Yuval Dor and Nish Patel for helpful discussions and advice. Work was supported by the Canadian Institutes of Health Research (grant FRN-343437 to R.K.) 6 and the National Institute of General Medical Sciences (grant GM26875 to M.K.), as well as a National Science Foundation Graduate Research Fellowship (M.B.G.). We also thank Patricia 8 and Alexander Younger and the Younger foundation for their generous donation to support our research.

